# Cdc25A phosphatase is activated and mediates neuronal cell death by PUMA via pRB/E2F1 pathway in a model of Parkinson’s disease

**DOI:** 10.1101/2024.05.06.592640

**Authors:** Anoy Kumar Das, Subhas C Biswas

## Abstract

Parkinson’s disease (PD) is a predominant movement disorder caused mainly due to selective loss of the dopaminergic neurons in the substantia nigra pars compacta of the mid brain. There is currently no cure for PD barring treatments to manage symptoms. The reasons might be due to lack of precise understanding of molecular mechanisms leading to neurodegeneration. Aberrant cell cycle activation has been implicated in neuronal death pathways of various neurodegenerative diseases including PD. This study investigates the role of cell cycle regulator Cell division cycle 25A (Cdc25A) in a PD-relevant neuron death model induced by 6-OHDA treatment. We find Cdc25A is rapidly elevated, activated and is playing a key role in neuron death by regulating Rb phosphorylation and E2F1 activity. Knockdown of Cdc25A via shRNA downregulates the levels of pro-apoptotic PUMA, an E2F1 target and cleaved Caspase-3 levels, suggesting Cdc25A may regulate neuronal apoptosis through these effectors. Our work sheds light on the intricate signaling networks involved in neurodegeneration and highlights Cdc25A as a potential therapeutic target for mitigating aberrant cell cycle re-entry underlying PD pathogenesis. These novel insights into molecular mechanisms provide a foundation for future development of neuroprotective strategies to slow or prevent progression of this debilitating disease.

## Introduction

Parkinson’s disease (PD) is a severe neurological ailment impacting millions of people worldwide. Projections indicate that by the year 2030, Parkinson’s disease (PD) will affect an estimated 8.67 million people globally (Calabrese 2007). A recent study (Willis *et al*. 2022) revealed a significant growth in PD incidence within the United States, with nearly 90,000 diagnoses in 2022 alone. This represents a 50% increase compared to previous estimates. Underlying cause of the disease is severe loss of the dopaminergic neurons in the substantia nigra pars compacta (SNpc) of the midbrain. These neurons are responsible for producing dopamine, which is necessary for basal ganglia mediated control of motor functions. Loss of dopamine results in the characteristic symptoms of PD, including tremors, rigidity, gait and difficulty in walking (Dauer & Przedborski 2003). The etiology of PD is uncertain, but hereditary and environmental factors are suspected. Although just a few genes have been found to be responsible for the familial type of the disease, the majority of cases of PD are sporadic (Papapetropoulos et al. 2007). While treatments are available to help manage its symptoms, there is still much work to be done in terms of finding a cure or prevention of the disease.

Neuronal cells have specialized functions, once differentiated, they remain in a quiescent state of cell cycle. Interestingly recent reports have implicated abnormal cell cycle re-entry (CCR) of differentiated neurons as one of the crucial steps observed in neurodegeneration and remain an intriguing subject of research (Joseph et al. 2020).

In Alzheimer’s disease (AD) patients, Cyclin-dependent kinase 4 (Cdk4) has been shown to be activated in the brain and is involved in apoptotic death induced by amyloid-β in cultured cortical neurons (Giovanni *et al*. 1999; Greene *et al*. 2004; Biswas *et al*. 2007). The Cdk4 protein is also induced in neurons that undergo apoptosis in AD (Busser et al. 1998; Ino & Chiba 2001), inhibiting Cdk4 protected neuronal PC12 cells from Amyloid-β mediated toxicity (Sanphui *et al*. 2013).

Evidence suggests CCR is also implicated in dopaminergic neuron death in PD models. The expression of cyclin D and E increases in cultured neurons after MPP+ (1-methyl-4-phenylpyridinium, a potent PD mimetic) exposure, indicating activation of Cdk4/6 and Cdk2 (Levy et al. 2009).

Among the cell cycle regulators implicated across these neurodegenerative disease models, the dual-specificity phosphatase Cell division cycle 25A (CDC25A) plays a key role in activating CDKs at the G1/S transition. Cdc25A is a dual-specificity phosphatase that cleaves phosphate groups from phosphorylated tyrosine and serine/threonine residues to activate Cdks activity at the G1-S transition (Aressy & Ducommun 2008). A variety of normal cells and tissues express Cdc25A during early embryonic development and in adults. In the postmitotic stage, cells that are destined to become neurons exit the cell cycle and enter a quiescent state. In proliferating cells, many genes that are vital for cycle completion are upregulated during the G1-S transition. Phosphatases remove inhibitory tyrosine/threonine phosphates from Cdks in order to activate cyclin-dependent kinases (e.g., Cdk4, Cdk2, etc.). Cdc25A is responsible for this crucial step during the G1-S transition (Nilsson & Hoffmann 2000). In recent studies, evidence suggests that Cdc25A level and activity increases during neuronal death induced by Camptothecin (Zhang et al. 2006), NGF deprivation, and beta-amyloid (Chatterjee et al. 2016).

6 hydroxydopamine (6-OHDA) has been utilized as PD mimetic to create a model to replicate cellular responses observed in PD, making it a powerful tool for investigating the molecular basis of cytotoxicity in PD (Hernandez-Baltazar et al. 2017; Cetin et al. 2022). This study examined the underlying mechanism connecting Cdc25A induction by 6-OHDA treatment with neuronal apoptosis.

## Materials and methods

### Material

NGF (cat no. N2513) and poly-d-lysine (cat no. P6407) were purchased from Sigma. Cell culture media Dulbecco’s modified Eagle’s medium (DMEM) (cat no. 31600-034), Neurobasal (cat no. 21103-049), B27, Pen Strep (cat no. 15140-122) Lipofectamine 2000 (cat no. PN 52887), Alexa Fluor (Cat no. A11008, A11003), horse serum (HS) (cat no. 26050088), and fetal bovine serum (FBS) (cat no.10082147) were purchased from Thermo Fisher Scientific. cDNA synthesis kit (cat no. 6110B) and real-time PCR kit (cat no. PN 4367659) were purchased from TAKARA. ECL reagent (cat no. RPN2232) and PVDF membrane (cat no. 10600023) were purchased from GE Healthcare. Anti-Cdc25A (cat no. SC-7389), anti-E2F1 (cat no. SC-251), anti-p-RB (cat no. sc-377528) and anti-Cdk4 (cat no. SC-260) antibodies were purchased from Santa Cruz Biotechnology technologies. Anti-PUMA (cat no. 4976S), anti-cleaved Caspase 3 (D175) (cat no. 9661S) were purchased from Cell Signaling Technology. Anti-TH antibodies (cat no. ab112) was purchased from Abcam. Primers were purchased from IDT DNA. Protein A-agarose (cat no. SC-2001) and Horseradish peroxidase-conjugated secondary antibodies (cat no. SC-2004; SC-2005) were from Santa Cruz Biotechnology.

### Cell Culture

Rat pheochromocytoma cells (PC12) (Research Resource Identifier: CVCL 0481) were grown as previously described (Greene & Tischler, 1976). Cells were maintained for 30 passages. Cells were maintained in DMEM medium containing 10% heat-inactivated HS and 5% heat-inactivated FBS, and neuronal differentiation was induced by NGF (50 ng/ml) for 7 days in DMEM medium containing 1% heat-inactivated HS. Experiments were performed on 5 DIV cells. Neuron-specific markers (immunostaining with MAP2 antibody) were validated before use to protect the cell line from cross-contamination.

Prior to treatment, primary dopaminergic (DA) neurons from the midbrain of an E-18 rat embryo were cultured in neurobasal media supplemented with B27 for 14 days (Weinert, Selvakumar, Tierney, & Alaviant, 2015). Penicillin-streptomycin was used as the antibiotic in both cases. Experiments were performed with 6-hydroxy-dopamine (6-OHDA) at a concentration of 100 µM in complete medium at different time points.

### Nuclear counting assay

Utilizing a detergent-containing buffer that dissolves the contents of the cell but leaves the nuclei intact, the intact nuclear counting test was carried out as described (Grau & Greene 2012). Then, a hemocytometer was used to count the intact nuclei. As a percentage of the overall cell population, the number of living cells was expressed.

### PCR

TRI reagent was used to isolate total RNA from the necessary samples (Sigma). Rat Cdc25A was amplified using the primer pairs 5′ CAGCTTCCACACCAGTCTCT-3′ and 5′-TTGACTGCCGATACCCATAT-3′. Both 5’-TCAACAGCAACTCCCACTCTT - 3’ and 5’ACCCTGTTGCTGTAGCCGTAT -3’ were used as GAPDH primers. Applied Bio systems 7500 Fast Real Time PCR System was used to run quantitative PCR utilizing One Step SYBR Ex Taq qRT-Takara while following the manufacturer’s instructions.

### Western Blot Analysis

PC12 cells that had undergone neuronal differentiation were lysed to extract proteins for analysis. As a method for protein analysis, Western blotting was utilized. In particular, 50 g of protein from each condition was separated on a 4-12% SDS-PAGE gel and then transferred to a PVDF membrane. The membrane was then probed with primary antibodies specific for the target proteins, and secondary antibodies conjugated with HRP were employed to detect the primary antibodies. Detection was performed using Biorad western blotting detection reagent, as per the manufacturer’s instructions. All Western blots were subsequently imaged using a geldoc (Azure) imaging system to capture the protein bands.

### Phosphatase Assay

The Cdc25A phosphatase test was conducted to assess the phosphatase enzyme Cdc25A’s activity. As an enzyme substrate, this assay employs the p-nitrophenyl phosphate liquid substrate system (obtained from Sigma). First, protein lysates were treated overnight at 4°C with anti-Cdc25A antibody while being shaken. The next day, 25 μl of protein A agarose was added to the mixture, which was then incubated for two hours. The Cdc25A coupled with agarose was then pelleted through centrifugation. The immunoprecipitated Cdc25A was then incubated at room temperature with 200 µl of p-nitro phenyl phosphate in the dark. Upon stopping the reaction, the absorbance of the solution was measured at 405 nm using an ELISA reader. This absorbance measurement is directly proportional to the Cdc25A phosphatase activity in the original protein lysate sample.

### Inhibitor Assay

Neuronally differentiated PC12 cells were co-treated with pharmacological inhibitors of Cdc25A, NSC95397 and 6-OHDA. Then the cells were kept overnight and counted using nuclear counting assay as described above.

### Immunocytochemistry

PC12 cells that had undergone neuronal differentiation were fixed with 4% paraformaldehyde and then blocked in blocking solution (3% NGS in 0.3% PBST) for two hours at room temperature. Primary antibodies were used to immune-label cells in a blocking solution and left on cells overnight at 4°C. The very following day, cells were washed in PBS before being incubated for 2 hours at room temperature with the appropriate secondary antibody. Hoechst was used to stain the nuclei.

### Transfection & Gene silencing

In order to silence the genes, the pSIREN-cdc25A-shRNA-ZsGreen and pSIREN-rand-shRNA-ZsGreen vectors have been used (Chatterjee *et al*. 2016). We transfected neuronal cells with 0.5 micrograms of plasmid per well in a total volume of 500 µl of serum-free DMEM using Lipofectamine 2000. A change of medium was followed by a gap of six hours, after which experiments were performed forty-eight hours after transfection.

### Survival Assay

A 24-well plate was used as the culture vessel for neuronal PC12 cells in this study. The cells were allowed to grow and proliferate for 3 days in the culture media before being subjected to transfection. The transfection was performed by introducing specific shRNAs into the cells using the Lipofectamine 2000 transfection reagent. After 48 hours of transfection, the cells were treated with 6-OHDA, a neurotoxin that is used to induce apoptosis of the transfected neurons.

To quantify the effectiveness of the transfection, the number of cells that remained transfected was counted using fluorescent microscopy. The transfected neurons were labeled with a green fluorescent marker and immediately (within 2 h) and after 24 and 48 hours of treatment, the number of transfected neurons (green cells) were counted under a fluorescence microscope.

### Animal care and housing

Male Sprague-Dawley rats (RRID: RGD_1566440), aged around 10 weeks and weighing 280-320g, were obtained from the animal facility of CSIR-Indian Institute of Chemical Biology, Kolkata. The animals were maintained on a 12 hours light and dark cycle, supplied food and water ad libitum and had humidity maintained at 60 ± 5%. The rats were sourced from the in-house breeding colony survived at CSIR-Indian Institute of Chemical Biology. The animals were kept in an adequately supervised environment, and in order to limit injury, two rats were positioned in ventilated cages during the treatment phase. The guidelines outlined by the Committee for the Purpose of Control and Supervision of Experiments on Animals under the Animal Welfare Divisions, Ministry of Environments and Forests, Government of India were followed during all the conducted experiments. The study was approved by the Institutional Animal Ethics Committee (reference number IICB/AEC-APP/MarchMeeting/2014). In the validation of hemiparkinsonian model, we used a total of 18 animals which was divided into two groups: the control group which was injected with vehicle (0.01% Ascorbic acid) and the experimental group which was injected with 6-OHDA. Each experiment was independently replicated three times, encompassing a total of twelve control animals and twelve animals treated with 6-OHDA. To check the *in vivo* level of Cdc25A, three animals were used for each experiment which was independently replicated for three times. This study did not utilize any sample size calculations or randomization procedures when assigning subjects to respective groups. Additionally, exclusion criteria were not explicitly predetermined for these experiments.

### Stereotaxic infusion

Anesthetization of the rats was carried out through the intraperitoneal injection of ketamine and xylazine at doses of 80 mg/kg and 8 mg/kg, respectively. The rats were then immobilized into a stereotaxic frame as the incisor bar was set 3.5 mm below the inter-aural line. A 6-OHDA infusion solution (4 μg in 1 μl), dissolved in 0.01% ascorbic acid, is then infused into the right MFB. The infusion aims at reaching the MFB accurately in rats through stereotaxic coordinates of anterio-posterior (AP) - 0.28 from the Bregma point, dorso-ventral (DV) - 0.82, and lateral (L) 0.15. (Haobam *et al*. 2015). Contralateral sides that received vehicle served as a control.

### Amphetamine- or apomorphine-induced rotations

Rotational behaviour was assessed to evaluate the Hemiparkinsonian model. The rats were placed in Perspex, transparent cylindrical chambers (45 cm diameter, 40 cm high). The number of complete body rotations made by each animal following drug administration was manually counted. Amphetamine (2.5 mg/kg, i.p.) was administered on the 14th day post-surgery, while apomorphine (0.1 mg/kg, subcutaneously) was administered on the 16th day post-surgery (Haobam *et al*. 2015).

### Immunohistochemistry of brain slices

Animals were anesthetized on the 16th or 8th day after toxin infusion (depending on the experimental setup) followed by perfusion. Initially, the rat brains were perfused using 0.1 M PBS. Next, the perfusion was performed using a 4% paraformaldehyde fixative solution to preserve tissue morphology. Subsequently, the brains were gently removed from the skull and left in the same 4% paraformaldehyde solution in a fridge overnight. After fixation, the brains were immersed in a 30% sucrose solution for cryoprotection. Finally, 20 μm sections spanning the substantia nigra pars compacta (SNpc) region and striatal regions were sliced using a cryostat and then embedded onto slides for further investigations. 20 μm rat brain cryosections, were blocked with 5% goat serum in PBS with 0.3% Triton-X 100 for 1 h at room temperature. Then the sections were incubated overnight at 4°C in primary antibody solution. Next the sections were washed three times with PBS, and secondary antibodies were incubated at room temperature for two hours. Following three washes with PBS and Hoechst dye nuclear staining, the sections were mounted and observed under fluorescence or confocal microscopy.

### Statistics

Experimental results are presented as Mean ± S.E.M. /S.D. The significance of differences between experimental and control group means is evaluated using the Student’s t-test, which uses unpaired, two-tailed arrays. The results are presented in the form of P-values. For datasets that contained more than two groups, one-way ANOVA was performed along with Tukey post-hoc test follow-up. All statistical calculation was done in Graphpad Prism 9.

## Results

### Cdc25A is upregulated in dopaminergic neurons *in vitro* and *in vivo* upon 6-OHDA treatment

In order to determine whether Cdc25A contributes to neuronal death in PD, we used neuronally differentiated PC12 cells that were treated with 6-OHDA. PC12 cells express the D2 isoform of the dopamine receptor (Zhu *et al*. 1997), which is responsive to 6-OHDA and has been employed in a number of PD studies (Sanphui *et al*. 2020). Western blotting analysis revealed that Cdc25A protein levels were increased significantly with time and as early as by 4h in neuronal PC12 cells in response to 6-OHDA treatment (Figure 1A,1B). Immunocytochemical analysis of neuronal PC12 cells also showed an increase in Cdc25A protein levels in response to 6-OHDA treatment (Figure 1C,1D). These experiments suggest that 6-OHDA-induced toxicity causes a rapid increase in Cdc25A protein levels in neuronal PC12 cells long before death occurs.

**Figure 1:**
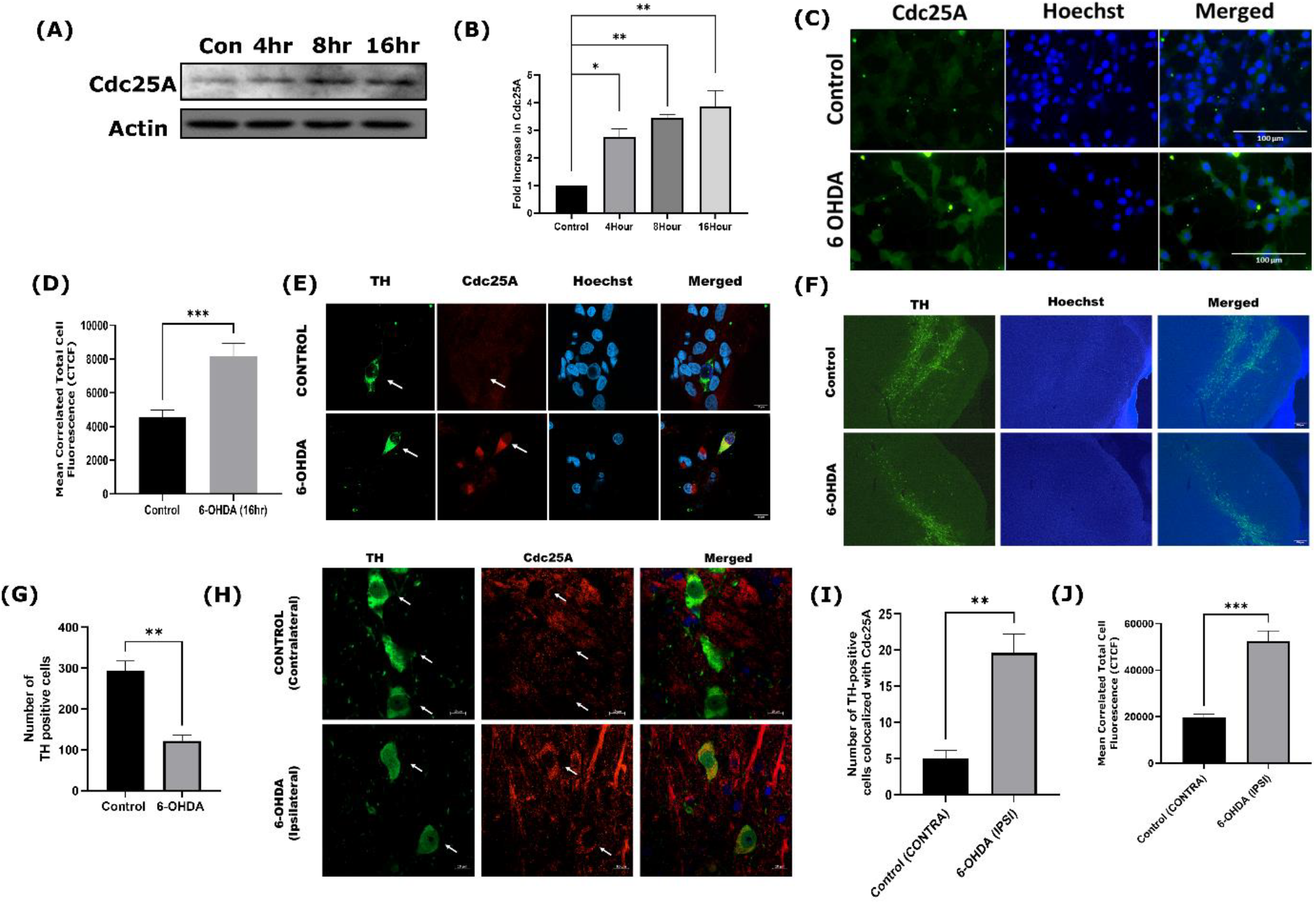
Increased expression of Cdc25A in response to 6-OHDA treatment *in vitro* and *in vivo*. (A) Neuronally differentiated PC12 cells were treated with 6-OHDA for indicated times. The total protein was subjected to western blotting analysis using Cdc25A antibody. *β*-Actin was used as a loading control. Representative immunoblot of increase in the levels of Cdc25A upon 6-OHDA treatment in neuronal PC12 cells. (B) Graphical representation of the Cdc25A protein levels as quantified by densitometry of western blots in neuronal PC12 cells subjected to 6-OHDA for 4h, 8 h, and 16 h respectively. Data are represented as mean ± S.E.M. of three independent experiments, ^*^*P*<0.05, ^**^p < 0.01. (C) Neuronally differentiated PC12 cells were treated with 6-OHDA for 16 hours and were processed for immunofluorescence. The representative image shows a significant increase in the Cdc25A level. (D) Quantitative analysis of the increase in Cdc25A in immunofluorescence level was calculated by Mean correlated total cell fluorescence (CTCF). Data represented was collected from three independent experiments. For each condition, 40 cells were counted per experiment. The asterisks denote statistically significant differences from the control at corresponding time points: ^***^p < 0.01. (E) A representative image of primary dopaminergic culture shows increases in the level of Cdc25A in TH+ cells (green). (F) Rats received unilateral injections of 6-OHDA or vehicle in the MFB at stereotaxic coordinates. Sixteen days post-surgery, the rats were euthanized and brain sections were immunostained with tyrosine hydroxylase (TH) for dopaminergic (DA) neurons (shown in green). 6-OHDA infused side had significantly fewer green TH-positive DA cells compared to that of control. A representative image is shown. (G) Graphical representation of number of TH-positive cells. ^**^p < 0. 01. (H) Representative images of co-immunostaining for TH marker protein and Cdc25A. (I) Graphical representation of number of cells having Cdc25A colocalized with TH positive cells: ^**^p < 0.01. (J) Quantitative analysis of increase in Cdc25A intensity in immunofluorescence level was calculated by mean correlated total cell fluorescence (CTCF). Data represented was collected from three independent experiment batches that consist of 3 animals in each batch. For each condition, 40 cells were counted per experiment. The asterisks denote statistically significant differences from control at corresponding time points: ^***^p < 0.001.

PD results in a selective loss of DA neurons in the SNpc of the midbrain. In order to determine whether Cdc25A is activated in DA neurons, we cultured primary DA neurons from rat brains as described before (Sanphui *et al*. 2020). Since the pure culture of SNpc DA neurons is practically impossible, we cultured DA neurons from the midbrain. Cdc25A levels were determined by immunocytochemistry in these primary cultured neurons. Cdc25A and TH (Tyrosine hydroxylase, DA neuron marker) antibodies were co-immunostained to identify the dopaminergic neurons. Results indicated a marked increase in Cdc25A levels in TH-positive neurons upon 6-OHDA treatment (Figure 1E).

To complement our culture studies, we next investigated whether Cdc25A is induced by 6-OHDA in *in vivo* models. Intracerebroventricular (ICV) administration of 6-hydroxydopamine (6-OHDA) is a well-established method to induce dopaminergic cell degeneration in animal models of PD (Rodriguez *et al*. 2001; Uretsky & Iversen 1970). In this study, we aimed to develop a hemiparkinsonian model in rats. The rats were divided into two groups and were unilaterally infused with 6-OHDA or vehicle targeting the medial forebrain bundle (MFB) using stereotaxic coordinates (as described in the Methods section). This approach selectively lesions the nigrostriatal dopaminergic pathway, mimicking a key feature of PD (Haobam *et al*. 2015). Post infusion the rats were subjected to amphetamine and apomorphine induced rotational behavior tests on day 14^th^ and 16^th^ post infusion respectively (as described in methods section) to validate the hemiparkinsonian model. Rotation test result confirmed the model (Supplementary Figure S1A-1D). For biochemical verification, the rats were sacrificed after 16 days and the brains were isolated, and after cryosectioning, the brain sections were processed for immunohistochemistry (IHC). The IHC data showed that the levels of TH-expressing cells drastically decreased on the 6-OHDA infused side (ipsilateral) as compared to the control side (Contralateral) (Figure 1F, 1G). Thus, the hemiparkinsonian model was validated. Next, we determined if the Cdc25A protein level increases *in vivo* in dopaminergic neurons. We infused 6-OHDA or vehicle (0.01% Ascorbic acid) on ipsilateral and contralateral sides of the rat brain respectively. Post infusion animals were kept for 8 days for partial degeneration of SNpc neurons. The animals were then sacrificed and brain sections of these animals were co-immunostained with TH (for dopaminergic neurons) and Cdc25A antibodies. IHC studies of the brain sections showed a significant increase in Cdc25A protein levels in TH-positive cells in the infused hemisphere (Ipsilateral). Overall, these results suggest that Cdc25A is upregulated in both *in vitro* and *in vivo* models of PD (Figure 1H-1J).

### Cdc25A protein level is increased due to its reduced Ubiquitin mediated degradation and it has increased phosphatase activity upon 6-OHDA treatment

Next, we tested whether the increase in the Cdc25A protein level was due to increase of its transcription or its post-translational modification. To this end, neuronal PC12 cells were treated with 6-OHDA for different time periods and then subjected to PCR. Since, no changes were found in the levels of Cdc25A transcripts in semi-quantitative PCR, quantitative RT-PCR was done and there were also no significant changes found (Figure 2A, 2B). While there was no significant change in Cdc25A mRNA levels in response to 6-OHDA, but Cdc25A protein levels did change as described above. We hypothesized that the Cdc25A protein is getting stabilized because of 6-OHDA treatment. Previous studies suggested that Cdc25A can be stabilized in the disease state due to decreased level of ubiquitination(Pereg *et al*. 2010). So, to check this hypothesis, coimmunoprecipitation of Cdc25A and Ubiquitin was performed by immunoprecipitation of Cdc25A followed by immunoblotting with Ubiquitin. Results showed that Cdc25A and Ubiquitin bind together and ubiquitination level of Cdc25A significantly goes down in 6-OHDA treated conditions (Figure 2C, 2D).

**Figure 2:**
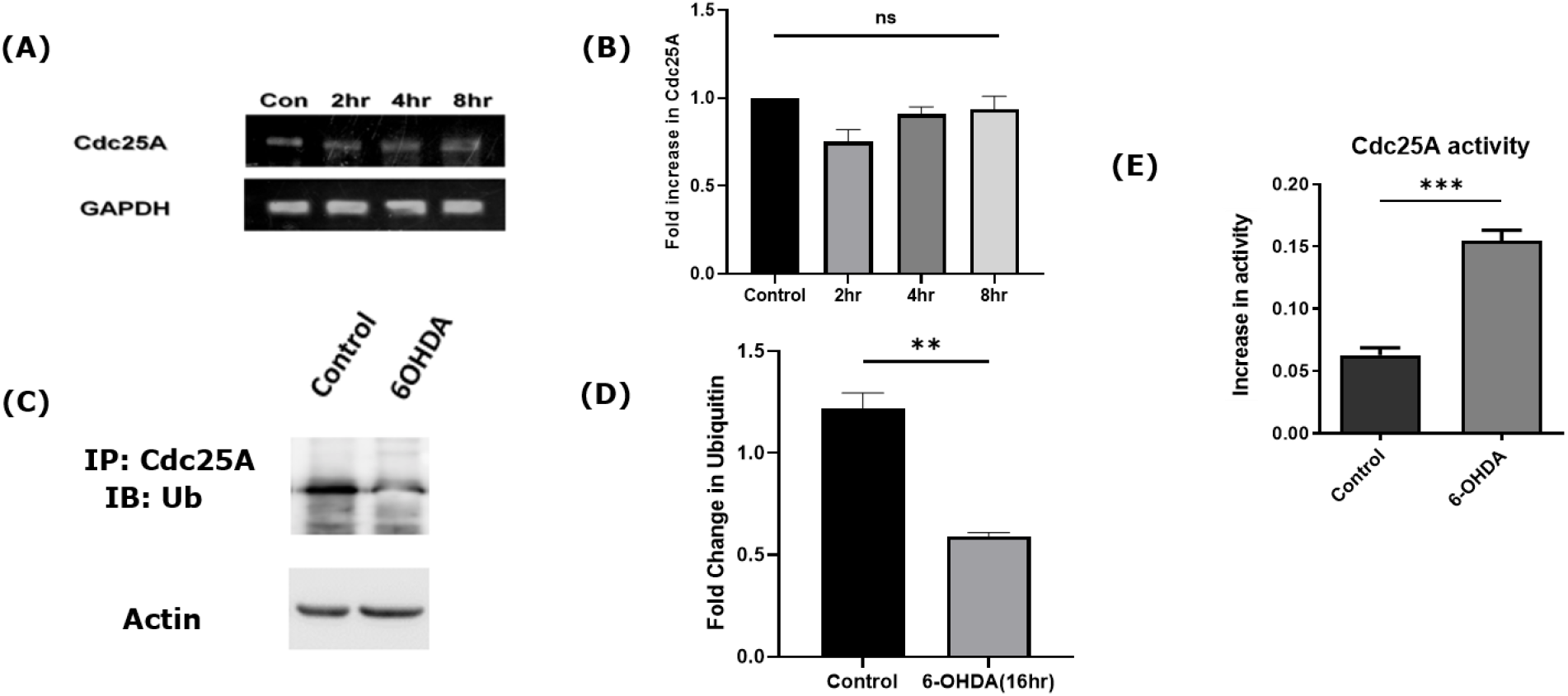
Cdc25A is stabilized and has increased phosphatase activity after 6-OHDA treatment. (A) Neuronally differentiated PC12 cells were subjected to 6-OHDA treatment for indicated times and total RNA was isolated from harvested cells. The RNA was analyzed by semiquantitative PCR for Cdc25A transcript levels. GAPDH was used as a loading control. Representative figure of three independent experiments with similar results is shown. (B) Quantitative real-time PCR of Cdc25A transcripts levels. GAPDH was used as a loading control. Data are represented as mean ± S.E.M. of four independent experiments. (C) Differentiated PC12 cells were treated with 6-OHDA for overnight and the cell lysates were immunoprecipitated (IP) with Cdc25A antibody and immunoblotted (IB) for Ubiquitin to check the change in the level of ubiquitination of Cdc25A upon 6-OHDA treatment. Representative image of three independent experiments with similar results is shown. (D) Quantitative analysis of fold change of Ubiquitin upon 6-OHDA treatment. Data are represented as mean ± S.E.M. of three independent experiments. ^**^p < 0.01 (E) Differentiated PC12 cells were subjected to 6-OHDA treatment for 8 h. Cdc25A was immunoprecipitated and subjected to phosphatase activity assay as described under ‘Materials and Methods’. Data represent mean ± S.E.M. of three independent experiments. The asterisks denote statistically significant differences from control at corresponding time points: ^***^p < 0.001.

We have also checked the activity of Cdc25A after 6-OHDA treatment. For this purpose, we performed a phosphatase activity assay because Cdc25A is a dual-activity phosphatase, and the results showed that the activity of Cdc25A is increased significantly after 6-OHDA treatment (Figure 2E). Overall, these experiments suggest that treatment of neuronal PC12 cells with 6-OHDA cause stabilization of Cdc25A protein and increases its activity long before cell death occurs.

### Downstream cell cycle molecules of Cdc25A are also induced in response to 6-OHDA treatment

Since Cdc25A acts as a major regulator of G1/S transition of the cell cycle pathway we determined the levels and activation of other cell cycle molecules of this phase of cell cycle in response to 6-OHDA as well. For that purpose, we treated the neuronally differentiated PC12 cells with 6-OHDA and checked the protein levels of Cdk4, pRB, and E2F1. While the levels of Cdk4 and pRB were increased in 16-hour time point, the level of E2F1 did not change significantly in response to 6-OHDA (Figure 3), however, its activity has increased (please see below). This indicates that other cell cycle molecules downstream of Cdc25A are also aberrantly activated in response to 6-OHDA.

**Figure 3:**
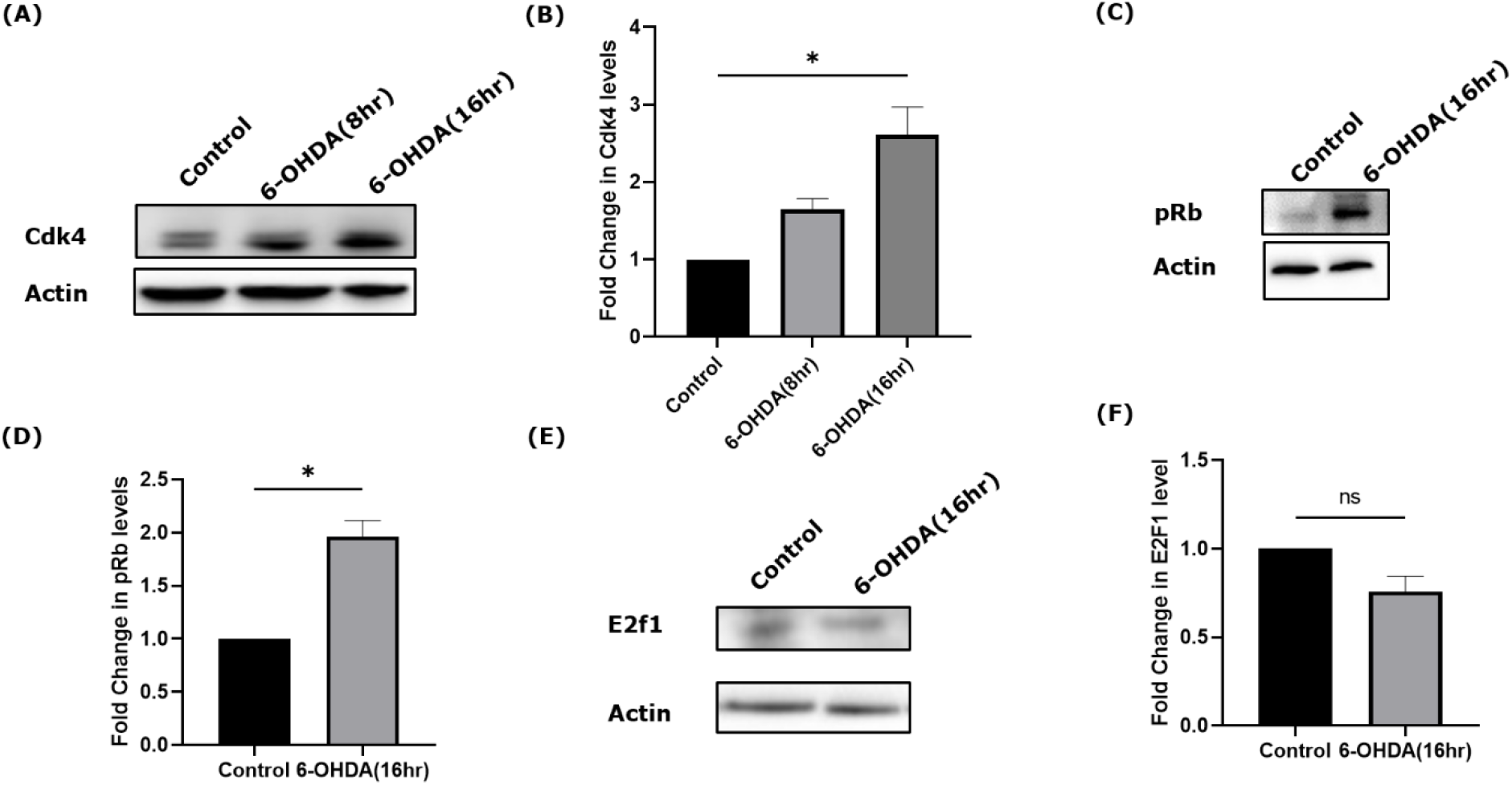
Cdk4 and pRB levels are increased in response to 6-OHDA treatment. Neuronally differentiated PC12 cells were treated with 6-OHDA for indicated times. Total protein was subjected to western blotting analysis to quantify levels of other cell cycle proteins. *β*-Actin was used as a loading control. Representative immunoblots of Cdk4, pRb and E2F1 upon 6-OHDA treatment in neuronal PC12 cells were presented on (A), (C) and (E) respectively. Quantitative analysis of the fold change of Cdk4, pRb and E2F1 was done by densitometry of western blots in neuronal PC12 cells and was presented on (B), (D) and (F) respectively. Data represent mean ± S.E.M. of three independent experiments. The asterisks denote statistically significant differences from control at corresponding time points: ^*^p < 0.05.

### Cdc25A is required for neuronal cell death caused by 6-OHDA treatment

Next, we investigated whether Cdc25A is required for 6-OHDA-induced neuronal cell death. To check this possibility, we treated neuronal PC12 cells with 6-OHDA in the presence and absence of the commercial Cdc25 inhibitor NSC95397 and examined cell viability. The drug protected the cells from degeneration (Figure 4A, 4B).

**Figure 4:**
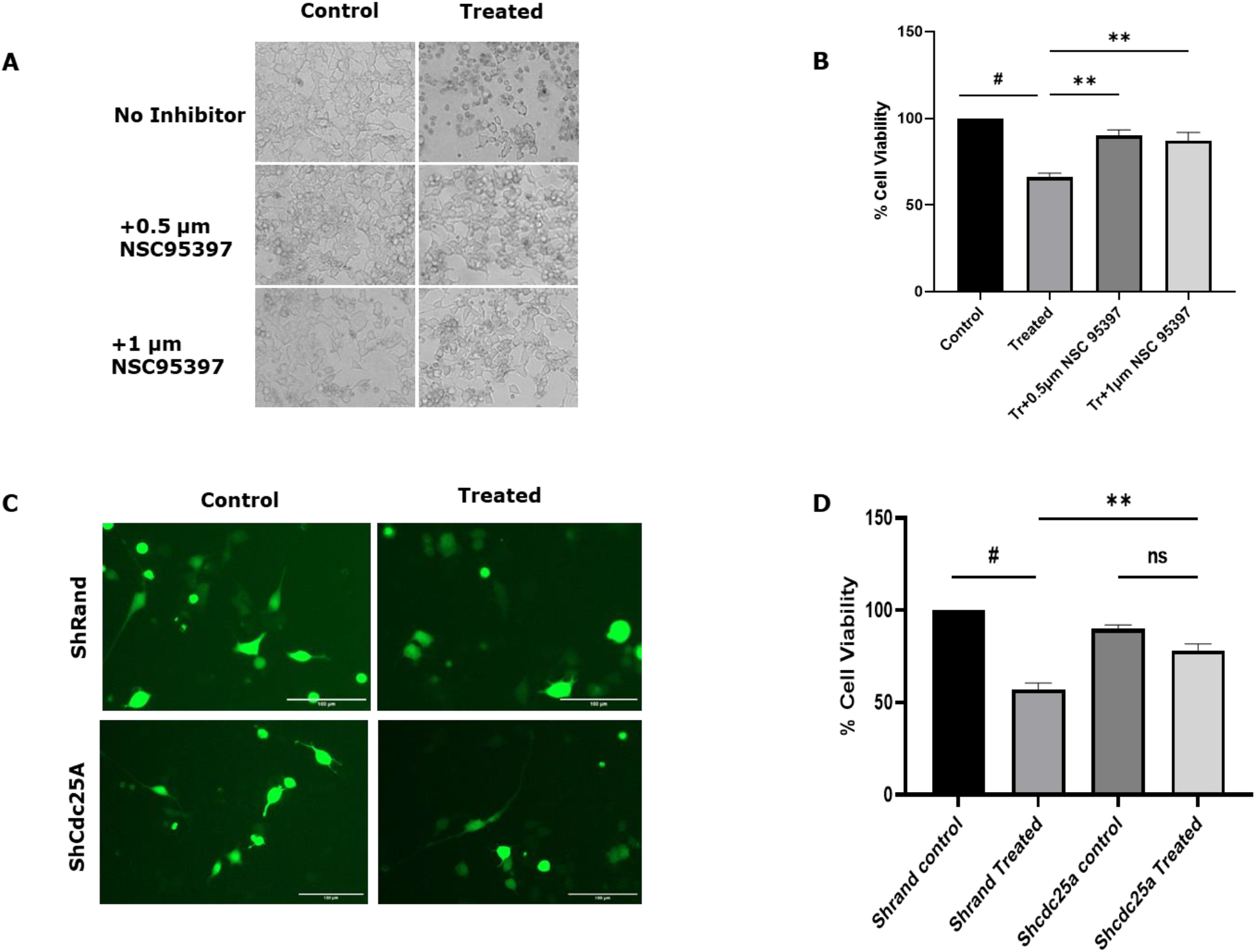
Cdc25 inhibition by a pan Cdc25 inhibitor and knockdown of Cdc25A prevent neuronal cell death in response to 6-OHDA. (A) Neuronal PC12 cells (5 DIV) were subjected to 6-OHDA in the presence and absence of NSC95397 at the indicated concentrations for 16 h. Representative phase contrast micrographs show retention of neurites after 6-OHDA treatment. (B) Graphical representation of the relative percentage of viable cells under the indicated conditions. Data are represented as mean ± S.E.M. of three independent experiments performed in duplicates. The asterisk denotes a statistically significant difference between indicated classes: #*P*<0.05, ^**^ *P*<0.01. (C) Differentiated PC12 cells (5 DIV) were transfected with shRand-zsgreen or shCdc25A-zsgreen and 48 h post-transfection, the cells were subjected to 6-OHDA treatment. The numbers of surviving-transfected (green) cells were counted at the indicated times under fluorescence microscopy. Representative images were shown in the panel. (D) Graphical Representation of percent viable cells of the survival experiment. Data are represented as mean ± SEM of three independent experiments. The asterisk denotes a statistically significant difference between indicated classes: #*P*<0.05. ^**^ *P*<0.01.

**Figure 5:**
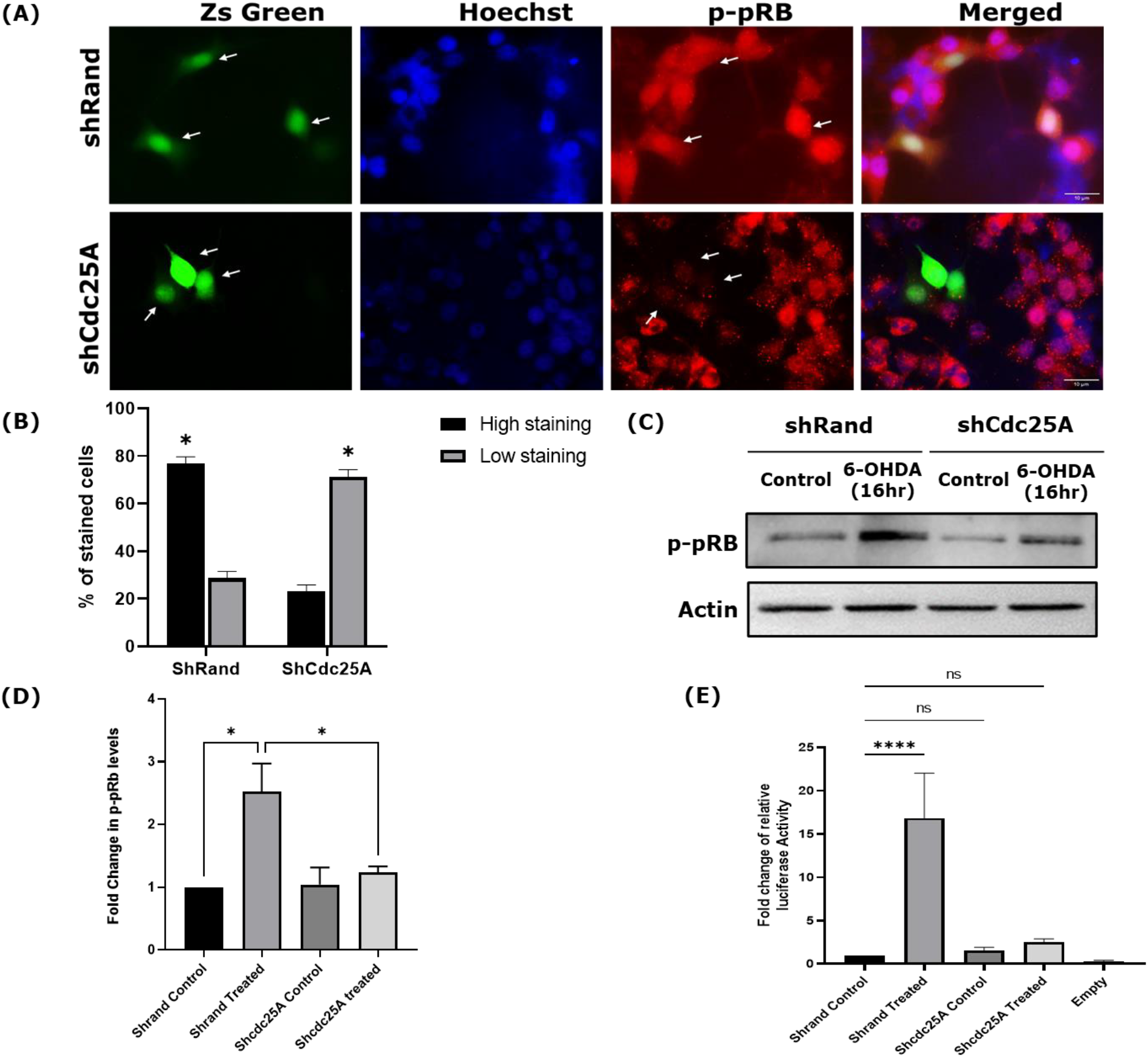
Knockdown of Cdc25A inhibits the phosphorylation of Rb and prevents the transcriptional activity of E2F1. (A) Differentiated PC12 cells (5 DIV) were transfected with shRand-zsgreen or shCdc25A-zsgreen and 48 h post transfection, the cells were subjected to 6-OHDA treatment and immunostained for phospho-pRb (p-pRb, red) and checked under a fluorescence microscope. (B) Quantitative representation of percentage level of p-pRb in transfected cells for indicated conditions. The asterisks denote statistically significant differences from control at corresponding time points: ^*^p < 0.05. (C) Differentiated PC12 cells (5 DIV) were transfected with shRand-zsgreen or shCdc25A-zsgreen and 48 h post transfection, the cells were subjected to 6-OHDA treatment. After 16 h incubation the cells were collected, lysed and processed for immunoblotting. Representative blot for p-pRb for indicated conditions is presented. (D) Graphical representation of the phosphorylation of Rb protein levels as quantified by densitometry of western blots for mentioned conditions. Data are represented as mean ± S.E.M. of three independent experiments, ^**^*P*<0.01. (E) Differentiated PC12 cells (5 DIV) were co-transfected with shRand-zsgreen or shCdc25A-zsgreen along with PUMA-LUCI construct and CMV-Renilla. 48 h post transfection, the cells were subjected to 6-OHDA treatment. After 16 h incubation the cells were collected and processed for Luciferase assay according to the manufacturer’s protocol (Promega). Quantification of relative Luciferase activity is represented graphically. The asterisks denote statistically significant differences from control at corresponding time points: ^*^p < 0.05, ^**^p < 0.01, ^****^p < 0.0001, n=3.

Because inhibitors can have nonspecific effects and are not selective, we used shRNA to specifically suppress the expression of Cdc25A. Compared with a control construct, shCdc25A significantly protected neuronal PC12 cells from death induced by 6-OHDA (Figure 4C, 4D). Altogether, these experiments suggest that Cdc25A plays a necessary role in neuronal death evoked by 6-OHDA.

### Cdc25A is required for Rb phosphorylation, E2F1 activation upon 6-OHDA treatment

Reports have shown that aberrant activation of the cell cycle pathway in neurons leads to neurodegeneration (Chatterjee *et al*. 2016; Greene *et al*. 2007). Reportedly, a E2F1 target, pro-apoptotic protein PUMA has shown to be upregulated in the PD models (Sanphui *et al*. 2020). Thus, we determined whether Cdc25A activation in turn phosphorylates Rb family proteins, triggering a series of events leading to E2F1 activation, induction of PUMA, caspase3 activation and subsequent neuronal apoptosis. If Cdc25A is upstream of the Rb proteins, then suppression/inhibition of Cdc25A should in turn block Rb phosphorylation and subsequent activation of caspase-3 by 6-OHDA treatment. When we examined PC12 cells treated with 6-OHDA by immunostaining, we found that shCdc25A, which knocked down the level of Cdc25A, greatly reduced the number of cells with high phospho-Rb levels. Blocking Cdc25A expression significantly suppressed the phosphorylation of Rb induced by 6-OHDA (Figure 6A-6D). Now phosphorylation of Rb should effectively activate the E2F1 transcription factor and it is known that E2F1 can activate the transcription of PUMA. So even though the levels of E2F1 were not altered we wanted to check the activity of E2F1. For that purpose, we used a luciferase-reporter probe that contained the PUMA promoter having E2F1 responsive elements, thus induction of the PUMA promoter by activated E2F1 will lead to the luciferase gene expression. Indeed, 6-OHDA treatment led to induction of PUMA promoter driven luciferase activity in control (shRand) transfected cells. In contrast, ShCdc25A was able to successfully block the E2F1-responsive PUMA promoter driven luciferase reporter gene expression (Figure 6E). Thus, this result suggests that knocking down the Cdc25A expression effectively repressed the activation of E2F1.

**Figure 6:**
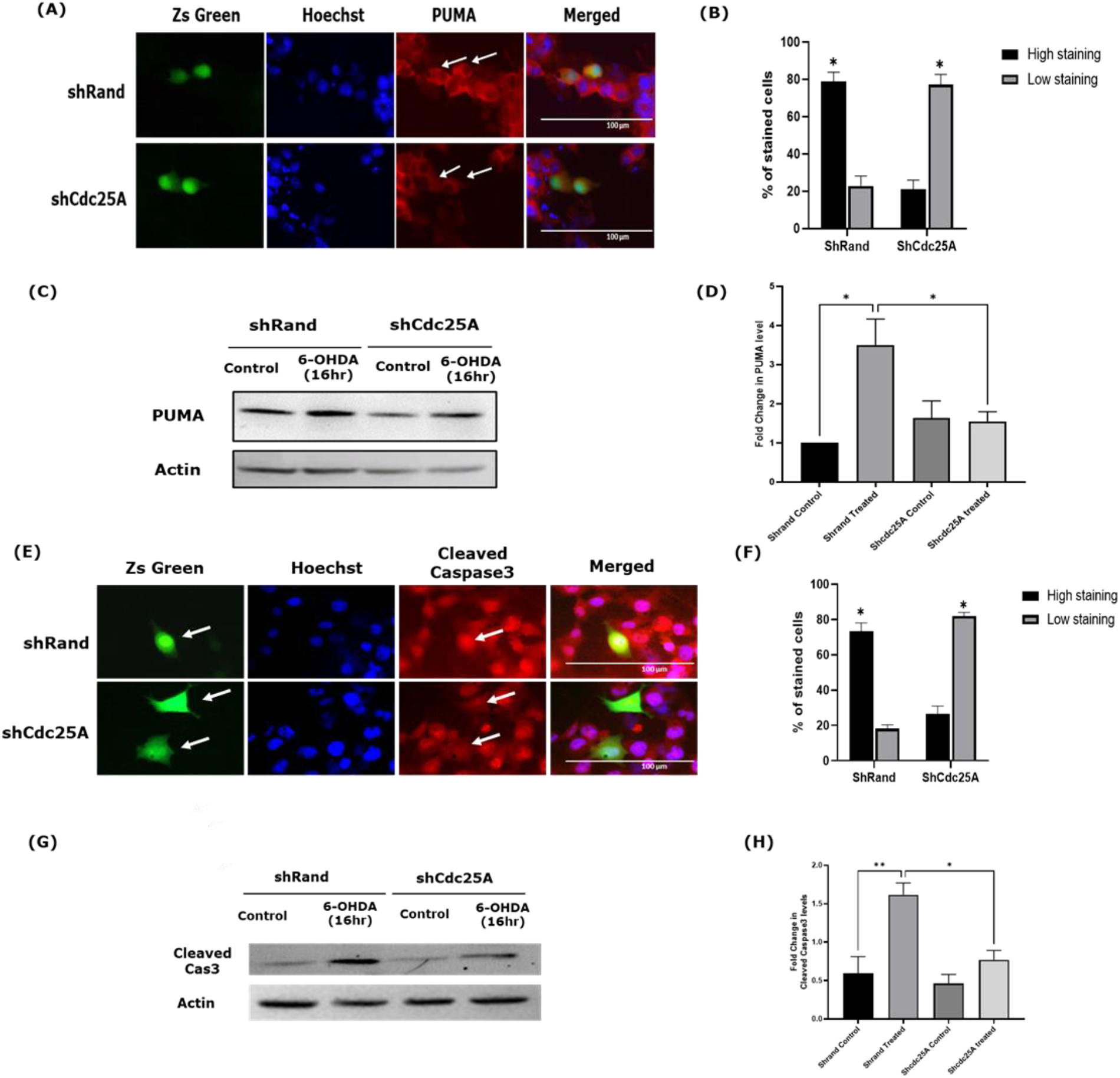
Knockdown of Cdc25A inhibits upregulation proapoptotic protein PUMA and cleaved Caspase3. (A) Differentiated PC12 cells (5 DIV) were transfected with shRand-zsgreen or shCdc25A-zsgreen and 48 h post transfection, the cells were subjected to 6-OHDA treatment and immunostained for PUMA (red) and checked under a fluorescence microscope. Representative image was shown. (B) Quantitative representation of percentage level of PUMA in transfected cells for indicated conditions. The asterisks denote statistically significant differences from control at corresponding time points: ^*^p < 0.05 (C) Differentiated PC12 cells (5 DIV) were transfected with shRand-zsgreen or shCdc25A-zsgreen and 48 h post transfection, the cells were subjected to 6-OHDA treatment. After 16 h incubation the cells were collected, lysed and processed for immunoblotting. Representative blot for PUMA for indicated conditions is presented. (D) Graphical representation of the PUMA protein levels as quantified by densitometry of western blots for mentioned conditions. Data are represented as mean ± S.E.M. of three independent experiments, ^*^*P*<0.05. (E) Differentiated PC12 cells (5 DIV) were transfected with shRand-zsgreen or shCdc25A-zsgreen and 48 h post transfection, the cells were subjected to 6-OHDA treatment and immunostained for Cleaved Caspase 3 (red) and checked under a fluorescence microscope. Representative image was shown. (F) Quantitative representation of percentage level of Cleaved Caspase 3 in transfected cells for indicated conditions. The asterisks denote statistically significant differences from control at corresponding time points: ^*^p < 0.05. (G) Differentiated PC12 cells (5 DIV) were transfected with shRand-zsgreen or shCdc25A-zsgreen and 48 h post transfection, the cells were subjected to 6-OHDA treatment. After 16 h incubation the cells were collected, lysed and processed for immunoblotting. Representative blot for Cleaved Caspase 3 for indicated conditions is presented. (H) Graphical representation of the Cleaved Caspase 3 protein levels as quantified by densitometry of western blots for mentioned conditions. Data are represented as mean ± S.E.M. of three independent experiments, ^*^*P*<0.05, ^**^p < 0.01.

### Inhibition of Cdc25A prevents induction of PUMA and caspase-3 cleavage upon 6-OHDA treatment

Finally, we checked the levels of PUMA and cleaved caspase-3 in Cdc25A knock-downed neuronal PC12 cells after 6-OHDA treatment to confirm the previous observation. We found that inhibition of Cdc25A also significantly blocked 6-OHDA-induced PUMA upregulation (Figure 6A-6D) and caspase-3 cleavage (Figure 6E-6H). Thus, these experiments suggest that Cdc25A is upstream of and required for the activation of cell cycle-related signaling pathways involved in neuronal cell death after 6-OHDA treatment.

## Discussion

A number of scientific disciplines are involved in studying PD, which is a complex and multifaceted syndrome. The role of cell cycle molecules in the development and progression of this debilitating disease is a particular interest of this study. The aberrant activation of cell cycle molecules has been identified as a contributing factor to neuronal apoptosis in several neurodegenerative diseases and represents a key mechanism promoting neuronal death.

Extensive research findings indicate that when post-mitotic neurons are exposed to various apoptotic conditions such as lack of trophic factors, DNA damage, exposure to Aβ, and oxidative stress, they leave the G_0_ state of the cell cycle and activate cell cycle proteins abnormally (Zhang *et al*. 2006; Chatterjee *et al*. 2016). This activation can be fatal to the neurons. One of the initial responses observed is the activation of G1/S cyclin-dependent kinases (Cdks), such as Cdk4 (Busser *et al*. 1998; Giovanni *et al*. 1999). Chatterjee et al. (Chatterjee *et al*. 2016) showed that Cdc25A is one of the major player that gets activated in Alzheimer’s disease. Moreover, it has been shown that Cdc25A serves as a crucial mediator of neuronal cell death resulting from Camptothecin mediated DNA damage in neuronal cells (Zhang *et al*. 2006) and ischemic insult (Iyirhiaro *et al*. 2017). We found that Cdc25A is induced and the activity is enhanced significantly in response to 6-OHDA treatment. Importantly, inhibition of Cdc25A by chemical inhibitors or shRNA protected neurons from 6-OHDA toxicity.

The retinoblastoma (pRb) family proteins gets phosphorylated in neurodegenerative disease conditions (Iyirhiaro *et al*. 2017; Chatterjee *et al*. 2016), leading to the dissociation of repressor complexes that include E2F and pRb proteins such as p130 (Greene *et al*. 2004; Greene *et al*. 2007). Subsequently, the de-repression of E2F-binding genes occurs which results in induction of E2F-responsive gene expression. The pRb/E2F pathway, which is involved in regulating the cell cycle, plays a critical role in the induction of cell death observed in PD (Hoglinger *et al*. 2007). Our work identified Cdc25A as a regulator of Rb phosphorylation and E2F1 activity in neuron death in response to 6-OHDA treatment. In line with a major role for Cdc25A in these events, shRNA targeting Cdc25A diminished phospho-Rb levels induced by 6-OHDA. Subsequently the E2F1 activity also got downregulated.

Loss of E2F-Rb complexes causes the expression of several genes such as the B- and C-myb transcription factors, which activate the proapoptotic protein Bim (Biswas *et al*. 2005). Bim promotes caspase activation, ultimately leading to neuron death (Biswas *et al*. 2007). E2F1 induces lot of proapoptotic BH3-only molecules during apoptosis (Hershko & Ginsberg 2004). The phenomenon of E2F1-induced apoptosis has been well documented to occur via either transcription-dependent or independent mechanisms, as previously reported (Azuma-Hara *et al*. 1999; Phillips *et al*. 1999). It is noteworthy that such occurrence can be impeded by the manifestation of the dominant negative DP-1 protein or pRb (Giovanni *et al*. 1999; Hsieh *et al*. 1997; Park *et al*. 2000). In addition, both pRb and E2F1 have been found to have significant regulatory roles in neuronal cell death during the developmental stage, as evidenced by the extensive cell loss within the central nervous system resulting from the loss of pRb function in transgenic mice (Jacks *et al*. 1992). E2F-1 induces melanoma cell apoptosis via p53 upregulated modulator of apoptosis (PUMA) (Hao *et al*. 2007). We have shown previously that PUMA also gets upregulated in PD (Sanphui *et al*. 2020). Interestingly in this work we observed that E2F-responsive PUMA promoter driven luciferase reporter activity as well as PUMA protein levels are enhanced in the PD model and shRNA that targets Cdc25A results in a reduction of both PUMA reporter activity and PUMA protein levels, concomitant with the inhibition of cleavage of Caspase-3, thus inhibiting cell from undergoing apoptosis. This finding is of significant interest, as it suggests that Cdc25A may play a pivotal role in the regulation of apoptotic signaling, specifically by impacting the expression of PUMA and Caspase-3.

Altogether, our observations indicate that Cdc25A is rapidly elevated, activated and plays a required role in PD relevant model of neuron death. Cdc25A acts as a regulator of Rb phosphorylation and E2F1 mediated PUMA-Caspase3 activation in neuron death in response to 6-OHDA treatment. These observations have important implications for our understanding of the intricate network of signaling pathways involved in the process of cell death, and may have potential therapeutic applications for diseases characterized by aberrant cell cycle activation that leads to neurodegeneration. Further studies aimed at targeting Cdc25A could lead to the development of new treatments that may prevent or slow the progression of these devastating neurodegenerative disorders. In a complementary study, we have recently shown that newly synthesized molecules that inhibit Cdc25A phosphatase activity block neuronal cell death caused by NGF deprivation and 6-OHDA (Pramanik *et al*. 2023). While more *in vivo* and clinical research is needed to fully understand the implications of our observations, these findings represent a promising avenue of research that may ultimately lead to new therapies for patients suffering from PD.

## Acknowledgments

The work was supported by one of the 12^th^ Five Year Plan Projects, miND (BSC0115) of CSIR, Govt. of India. We would like to thank Dr. Paidi Ramesh Kumar for helping in stereotaxic surgery in animals.

## References

Azuma-Hara, M., Taniura, H., Uetsuki, T., Niinobe, M. and Yoshikawa, K. (1999) Regulation and deregulation of E2F1 in postmitotic neurons differentiated from embryonal carcinoma P19 cells. Exp Cell Res 251, 442–451.

Biswas, S. C., Liu, D. X. and Greene, L. A. (2005) Bim is a direct target of a neuronal E2F-dependent apoptotic pathway. J Neurosci 25, 8349–8358.

Biswas, S. C., Shi, Y., Sproul, A. and Greene, L. A. (2007) Pro-apoptotic Bim induction in response to nerve growth factor deprivation requires simultaneous activation of three different death signaling pathways. J Biol Chem 282, 29368–29374.

Busser, J., Geldmacher, D. S. and Herrup, K. (1998) Ectopic cell cycle proteins predict the sites of neuronal cell death in Alzheimer’s disease brain. J Neurosci 18, 2801–2807.

Calabrese, V. P. (2007) Projected number of people with Parkinson disease in the most populous nations, 2005 through 2030. Neurology 69, 223–224; author reply 224.

Chatterjee, N., Sanphui, P., Kemeny, S., Greene, L. A. and Biswas, S. C. (2016) Role and regulation of Cdc25A phosphatase in neuron death induced by NGF deprivation or beta-amyloid. Cell Death Discov 2, 16083.

Giovanni, A., Wirtz-Brugger, F., Keramaris, E., Slack, R. and Park, D. S. (1999) Involvement of cell cycle elements, cyclin-dependent kinases, pRb, and E2F x DP, in B-amyloid-induced neuronal death. J Biol Chem 274, 19011–19016.

Grau, C. M. and Greene, L. A. (2012) Use of PC12 cells and rat superior cervical ganglion sympathetic neurons as models for neuroprotective assays relevant to Parkinson’s disease. Methods Mol Biol 846, 201–211.

Greene, L. A., Biswas, S. C. and Liu, D. X. (2004) Cell cycle molecules and vertebrate neuron death: E2F at the hub. Cell Death Differ 11, 49–60.

Greene, L. A., Liu, D. X., Troy, C. M. and Biswas, S. C. (2007) Cell cycle molecules define a pathway required for neuron death in development and disease. Biochim Biophys Acta 1772, 392–401.

Hao, H., Dong, Y., Bowling, M. T., Gomez-Gutierrez, J. G., Zhou, H. S. and McMasters, K. M. (2007) E2F-1 induces melanoma cell apoptosis via PUMA up-regulation and Bax translocation. BMC Cancer 7, 24.

Haobam, R., Tripathy, D., Kaidery, N. A. and Mohanakumar, K. P. (2015) Embryonic stem cells derived neuron transplantation recovery in models of parkinsonism in relation to severity of the disorder in rats. Rejuvenation Res 18, 173–184.

Hershko, T. and Ginsberg, D. (2004) Up-regulation of Bcl-2 homology 3 (BH3)-only proteins by E2F1 mediates apoptosis. J Biol Chem 279, 8627–8634.

Hoglinger, G. U., Breunig, J. J., Depboylu, C. et al. (2007) The pRb/E2F cell-cycle pathway mediates cell death in Parkinson’s disease. Proc Natl Acad Sci U S A 104, 3585–3590.

Hsieh, J. K., Fredersdorf, S., Kouzarides, T., Martin, K. and Lu, X. (1997) E2F1-induced apoptosis requires DNA binding but not transactivation and is inhibited by the retinoblastoma protein through direct interaction. Genes Dev 11, 1840–1852.

Iyirhiaro, G. O., Im, D. S., Boonying, W., Callaghan, S. M., During, M. J., Slack, R. S. and Park, D. S. (2017) Cdc25A Is a Critical Mediator of Ischemic Neuronal Death In Vitro and In Vivo. J Neurosci 37, 6729–6740.

Jacks, T., Fazeli, A., Schmitt, E. M., Bronson, R. T., Goodell, M. A. and Weinberg, R. A. (1992) Effects of an Rb mutation in the mouse. Nature 359, 295–300.

Park, D. S., Morris, E. J., Bremner, R., Keramaris, E., Padmanabhan, J., Rosenbaum, M., Shelanski, M. L., Geller, H. M. and Greene, L. A. (2000) Involvement of retinoblastoma family members and E2F/DP complexes in the death of neurons evoked by DNA damage. J Neurosci 20, 3104–3114.

Pereg, Y., Liu, B. Y., O’Rourke, K. M., Sagolla, M., Dey, A., Komuves, L., French, D. M. and Dixit, V. M. (2010) Ubiquitin hydrolase Dub3 promotes oncogenic transformation by stabilizing Cdc25A. Nat Cell Biol 12, 400–406.

Phillips, A. C., Ernst, M. K., Bates, S., Rice, N. R. and Vousden, K. H. (1999) E2F-1 potentiates cell death by blocking antiapoptotic signaling pathways. Mol Cell 4, 771–781.

Pramanik, S. K., Sanphui, P., Das, A. K., Banerji, B. and Biswas, S. C. (2023) Small-Molecule Cdc25A Inhibitors Protect Neuronal Cells from Death Evoked by NGF Deprivation and 6-Hydroxydopamine. ACS Chem Neurosci 14, 1226–1237.

Rodriguez, M., Barroso-Chinea, P., Abdala, P., Obeso, J. and Gonzalez-Hernandez, T. (2001) Dopamine cell degeneration induced by intraventricular administration of 6-hydroxydopamine in the rat: similarities with cell loss in parkinson’s disease. Exp Neurol 169, 163–181.

Sanphui, P., Kumar Das, A. and Biswas, S. C. (2020) Forkhead Box O3a requires BAF57, a subunit of chromatin remodeler SWI/SNF complex for induction of p53 up-regulated modulator of apoptosis (Puma) in a model of Parkinson’s disease. J Neurochem 154, 547–561.

Sanphui, P., Pramanik, S. K., Chatterjee, N., Moorthi, P., Banerji, B. and Biswas, S. C. (2013) Efficacy of cyclin dependent kinase 4 inhibitors as potent neuroprotective agents against insults relevant to Alzheimer’s disease. PLoS One 8, e78842.

Uretsky, N. J. and Iversen, L. L. (1970) Effects of 6-hydroxydopamine on catecholamine containing neurones in the rat brain. J Neurochem 17, 269–278.

Willis, A. W., Roberts, E., Beck, J. C. et al. (2022) Incidence of Parkinson disease in North America. NPJ Parkinsons Dis 8, 170.

Zhang, Y., Qu, D., Morris, E. J., O’Hare, M. J., Callaghan, S. M., Slack, R. S., Geller, H. M. and Park, D. S. (2006) The Chk1/Cdc25A pathway as activators of the cell cycle in neuronal death induced by camptothecin. J Neurosci 26, 8819–8828.

Zhu, W. H., Conforti, L. and Millhorn, D. E. (1997) Expression of dopamine D2 receptor in PC-12 cells and regulation of membrane conductances by dopamine. Am J Physiol 273, C1143–1150.

