## Supplementary Figures for "Cdc25A phosphatase is activated and mediates neuronal cell death by PUMA via pRB/E2F1 pathway in a model of Parkinson’s disease"

**Supplementary Data**

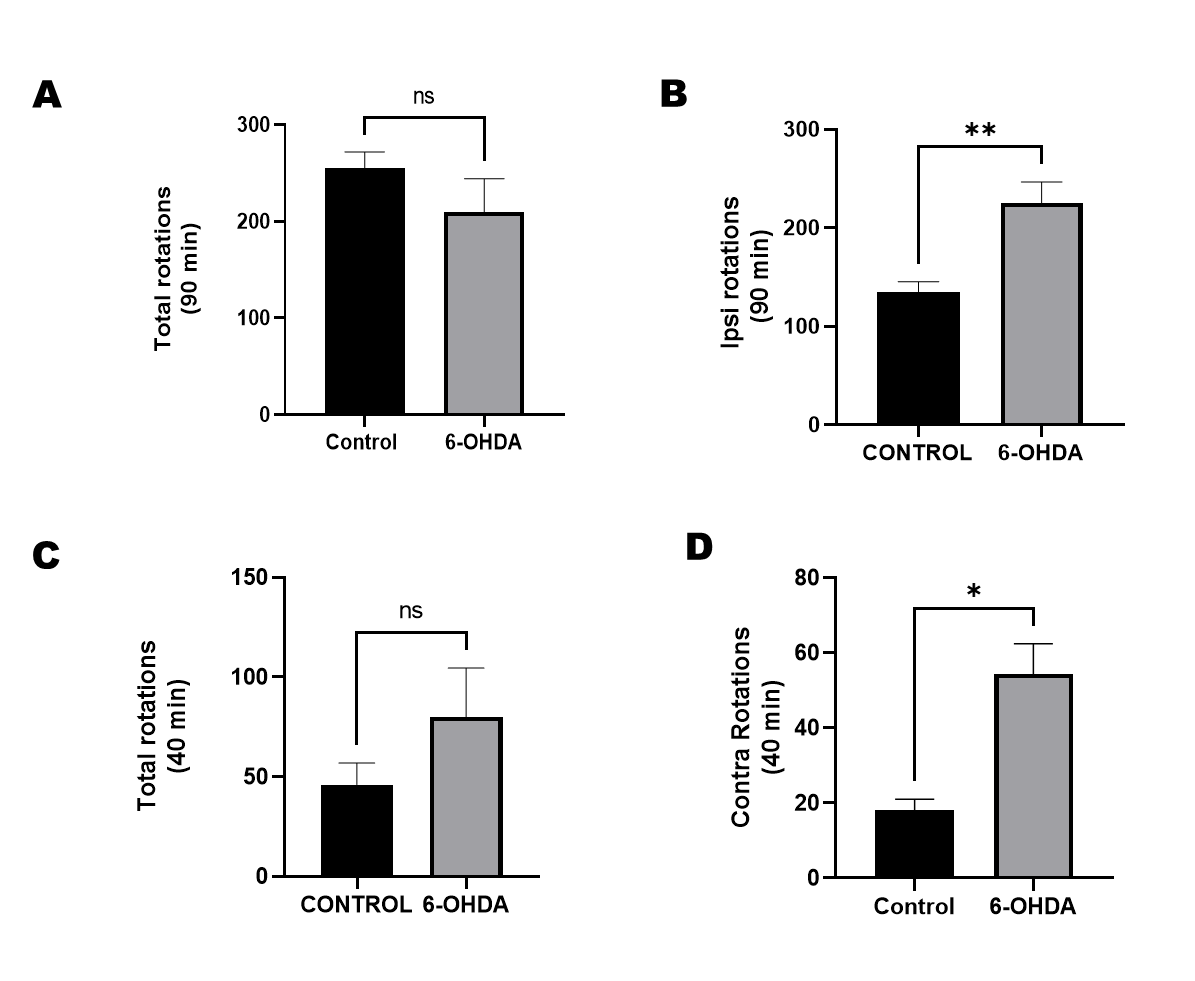

**Figure S1:** Behavioural effects of unilateral intra-median forebrain bundle (MFB) infusion of different doses of 6-hydroxydopamine (6-OHDA)

Rats that had undergone stereotactic infusion were subjected to rotation tests using amphetamine and apomorphine for 14 and 16 days, respectively. (A) and (B) graphical representation displays the changes in total rotations as well as Ipsi rotations after injecting amphetamine. Ipsi rotations increases significantly in case of 6-OHDA infused animals under the effect of amphetamine. The data represent the mean ± S.E.M. from three independent experiments, each consisting of four control animals and four toxin-infused animals per batch. Statistical significance is indicated by **P<0.01. While (C) and (D) graphical representation displays the changes in total rotations as well as contra rotations after injecting apomorphine. Contra rotations increases significantly in case of 6-OHDA infused animals under the effect of apomorphine. The data represent the mean ± S.E.M. from three independent experiments, each consisting of four control animals and four toxin-infused animals per batch. Statistical significance is indicated by *P<0.05
